# Effect of Elbow Flexion Angle on Cortical Voluntary Activation of the Non-Impaired Biceps: A 3 Days Repeated Sessions Study

**DOI:** 10.1101/2020.06.12.148536

**Authors:** Thibault Roumengous, Paul A. Howell, Carrie L. Peterson

## Abstract

Measurement of cortical voluntary activation (VA) with transcranial magnetic stimulation (TMS) is limited by technical challenges. One challenge is the difficulty in preferential stimulation of cortical neurons projecting to the target muscle and minimal stimulation of cortical neurons projecting to antagonists. Thus, the motor evoked potential (MEP) response to TMS in the target muscle compared to its primary antagonist may be an important parameter in the assessment of cortical VA. Modulating isometric elbow angle alters the magnitude of MEPs at rest. The purpose of this study was to evaluate the effect of isometric elbow flexion-extension angle on: 1) the ratio of biceps MEP relative to the triceps MEP amplitude across a range of voluntary efforts, and 2) cortical VA. Ten non-impaired participants completed three sessions wherein VA was determined using TMS at 45°, 90° and 120° of isometric elbow flexion, and peripheral electrical stimulation at 90° of elbow flexion. The biceps/triceps MEP ratio was greater in the more flexed elbow angle (120° flexion) compared to 90° during contractions of 50% and 75% of maximum voluntary contraction. Cortical VA assessed in the more extended elbow angle (45° flexion) was lower relative to 90° elbow flexion; this effect was dependent on the biceps/triceps MEP ratio. Cortical VA was sensitive to small changes in the linearity of the voluntary torque and superimposed twitch relationship, regardless of the elbow angle. Peripheral and cortical VA measures at 90° of elbow flexion were repeatable across three days. In conclusion, although the biceps/triceps MEP ratio was increased at a more flexed elbow angle relative to 90°, there was not a corresponding difference in cortical VA. Thus, increasing the MEP ratio via elbow angle did not affect estimation of cortical VA.

## INTRODUCTION

Voluntary activation is a measure which quantifies the level of voluntary drive to innervate muscle. Assessment of voluntary activation is useful in the study of neuromuscular impairments in clinical populations (1–3) and mechanisms of neuromuscular fatigue (4–6). Measurement of voluntary activation involves superimposing electrical stimulation of motoneurons upon an individual’s voluntary effort to activate muscle. A deficit in voluntary activation is indicated when muscle force during maximum voluntary effort is further increased by electrical stimulation. The increase in muscle force indicates that the stimulus recruited additional motor units beyond those already recruited via voluntary effort. When the stimulus is applied to peripheral motor nerve, we can assess what we will refer to as peripheral voluntary activation (peripheral VA). Peripheral VA indicates the net voluntary drive, consisting of cortical drive and the transmission of the neural signal through corticospinal and lower motoneurons to innervate muscle. Peripheral VA cannot indicate the site of deficit in voluntary drive within the pathway from cortex to muscle. Thus, a technique to assess cortical voluntary activation (cortical VA) via transcranial magnetic stimulation (TMS) was developed to indicate deficits in voluntary cortical drive (4). Together, measurement of cortical and peripheral VA can better localize the deficit in voluntary drive, which can be useful in directing rehabilitation strategies in patient populations.

Measurement of cortical VA is limited by technical challenges (7–9). Key technical challenges are that TMS over the motor cortex may: 1) activate cortical neurons projecting to muscles other than the target muscle, including antagonists, and 2) not activate every motor unit in the target muscle. An ideal measure of cortical VA would be obtained when TMS activates neurons that recruit all motor units not already recruited in the target muscle during maximum voluntary effort, and does not recruit antagonist motor units. The ratio of the target muscle motor evoked potential (MEP) in response to TMS relative to the antagonist MEP (i.e., target MEP amplitude divided by antagonist MEP amplitude) can indicate how well the ideal measurement scenario is achieved. In the assessment of cortical VA in non-impaired individuals, adjustment of the TMS pulse intensity and selection of an appropriate target muscle can optimize the target/antagonist MEP ratio (4). For example, when the biceps brachii is the target muscle for assessment of cortical VA in non-impaired individuals, stimulus intensity can be adjusted to elicit a biceps MEP amplitude that is greater than 50% of the maximal M-wave (Mmax), and a triceps MEP less than 20% Mmax (i.e., a biceps/triceps MEP ratio greater than 2.5) (7). However, experimental design may further optimize the target/antagonist MEP ratio, which may be important in the assessment of cortical VA in patient populations in which measurement of cortical VA has proved challenging (10).

Static changes in joint angle modulate MEP amplitudes in relaxed muscle (11,12). Therefore, careful prescription of the isometric joint angle is a promising approach to optimize the target/antagonist MEP ratio and improve the measurement of cortical VA. In previous work assessing relaxed muscle, biceps MEPs were maximized and triceps MEPs were minimized at a more flexed elbow angle relative to more extended elbow angles (12,13). However, whether joint angle can be prescribed to optimize the target/antagonist MEP ratio across the range of voluntary effort levels needed to estimate cortical VA remains unknown. In the current study we focus on modulation of the isometric elbow flexion-extension angle in the assessment of cortical VA of the biceps in non-impaired individuals. The primary objectives of this study were to determine the effect of the isometric elbow angle on: 1) the biceps/triceps MEP ratio across a range of voluntary efforts, and 2) cortical VA. We hypothesized that the biceps/triceps MEP ratio would be greatest in a more flexed elbow angle at each level of voluntary effort. Further, we hypothesized that cortical VA would depend on the biceps/triceps MEP ratio, based on our expectation that a greater biceps/triceps MEP ratio would indicate better targeting of the biceps relative to the triceps with TMS. A secondary objective of this study was to determine the repeatability of cortical and peripheral VA estimates across three sessions when measured at 90° of elbow flexion.

## METHODS

### Experiment Overview

Ten non-impaired individuals participated in three sessions (four females, six males, average age 22.7 ± 2.5 years). Participants were screened to ensure that they were eligible to receive TMS and provided informed written consent. The study was approved by the Institutional Review Board of Virginia Commonwealth University. In each session, participants completed trials to assess cortical and peripheral VA. Three sessions were conducted in order to assess the repeatability of cortical and peripheral VA in a common elbow angle (90°, elbow flexion-extension angle defined in accordance with recommendations of the International Society of Biomechanics (14). Sessions were separated by at least one day, with no more than 7 days between sessions. Each session consisted of up to three experimental blocks; one block of trials to assess peripheral VA with the elbow positioned at 90° of flexion, one block to assess cortical VA in 90° elbow flexion, and one block to assess cortical VA in either 45° or 120° elbow flexion. Cortical VA was not assessed at 45° and 120° in each session to minimize fatigue. The order of the blocks was randomized. Before each block, three maximum voluntary contractions (MVCs) were recorded at each isometric elbow angle. For all trials, the participant’s forearm was immobilized in a custom brace attached to a six degree-of-freedom load cell (Model 30E15A4-I40-EF-100L, JR3, Woodland, CA) (Figure 1). Force and moment data were sampled at 2000 Hz. EMG data were also sampled at 2000 Hz and bandwidth limited to 20-450 Hz.

**Figure 1.**
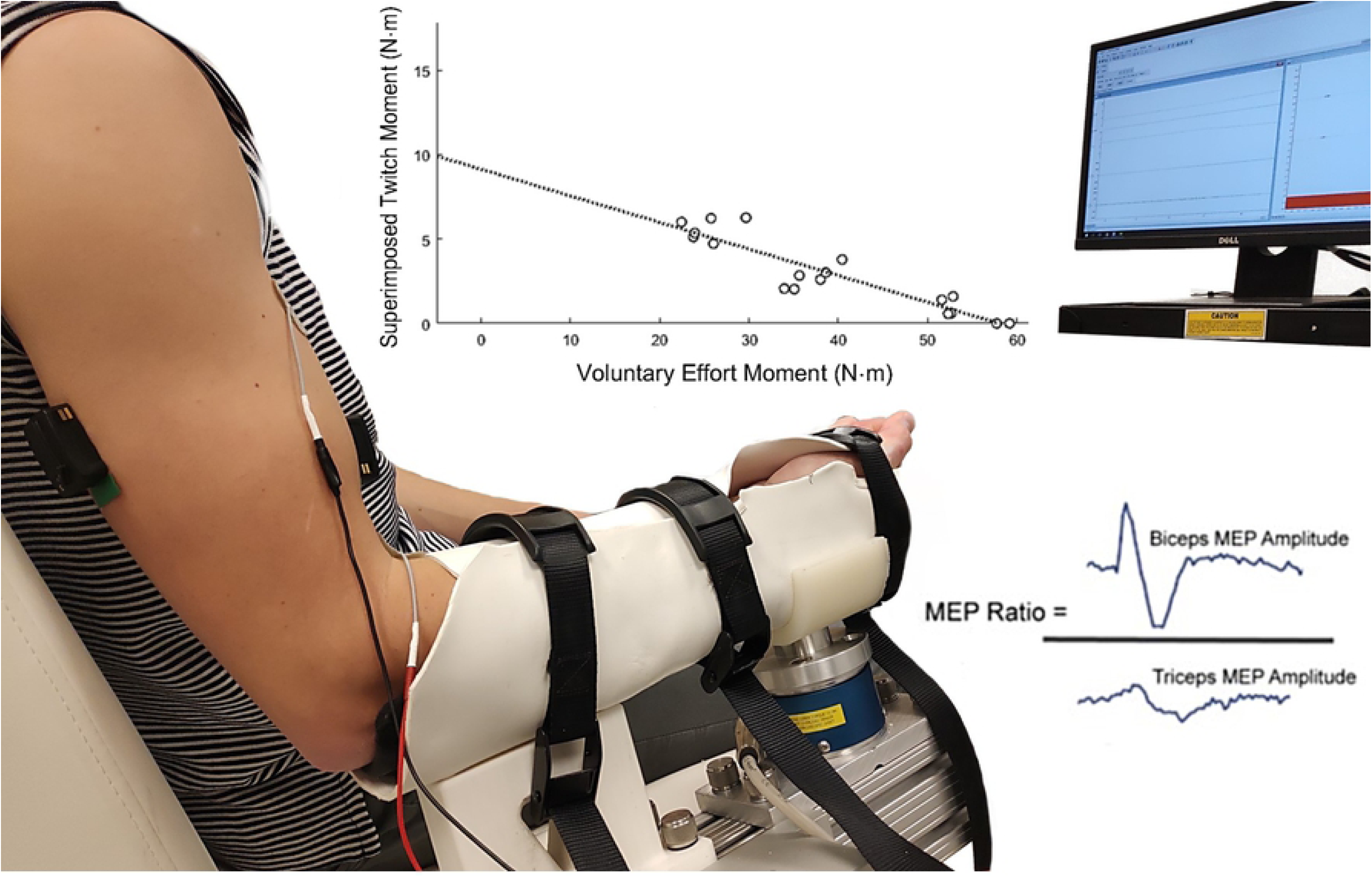
Experimental set up and moment and EMG signal recording paradigm. Participants received visual feedback of their voluntary moment as a thermometer-like gauge. Cortical VA trials were used to estimate the resting twitch via linear regression of superimposed twitch moments relative to the voluntary moments. Biceps and triceps EMG data were used to calculate the MEP ratio for each trial.

### Maximal Voluntary Contractions

Participants performed three MVCs of the elbow flexors for six seconds while receiving real-time visual moment feedback and verbal encouragement. Each maximum effort was separated by at least 90 seconds of rest. The participant’s MVC was calculated for each effort as the mean elbow moment maintained over ± 250 ms from the maximal moment achieved. The mean elbow flexion moment of three MVC trials was used for subsequent trials during which participants were asked to generate a voluntary moment to match a percentage of their MVC moment.

### Assessment of Peripheral VA

Participants completed trials during which motor point electrical stimulation was superimposed on isometric MVCs in elbow flexion in order to estimate peripheral VA. For motor point stimulation, stimulating electrodes were placed over the biceps belly and distal tendon. Stimulus intensity was determined by increasing the stimulation current in 10 mA increments until the moment response in the resting biceps reached a plateau. The threshold current (i.e., current corresponding to the start of the moment plateau) was recorded and motor point stimulation intensity was set at 130% of the threshold current. Using visual moment feedback, participants were instructed to perform nine MVCs in elbow flexion during which stimulation was superimposed during and after the voluntary effort. Motor point stimulation with a 0.2 ms pulse width (DS7AH, Digitimer, UK) was delivered after the participant maintained a voluntary moment ≥ 95% of their MVC moment for 0.5 seconds. A second stimulus event (same intensity and pulse width) was delivered six seconds after the first stimulus event while the arm was at rest.

### Assessment of Cortical VA

Single pulse TMS was delivered using a 126 mm diameter double cone coil (Magstim coil model) and Magstim BiStim^2^. Motor cortex mapping was performed each session to obtain the location that evoked the largest peak-to-peak MEP in the biceps relative to the triceps at the lowest stimulation intensity (15). This location was then marked on a cap secured to the participant’s head; subsequent stimuli were delivered at that location. Resting motor threshold (RMT) was determined as the lowest stimulus intensity able to induce biceps MEPs ≥ 50 µV in at least 5 out of 10 stimuli (16,17). In each session, participants completed a cortical VA block with their elbow flexed at 90°. Participants also completed a block in which isometric arm posture was modified to be either 45° or 120° of elbow flexion. Modified elbow flexion conditions were presented in a randomized order. Each block consisted of a set of 24 isometric contractions of the elbow flexors in randomized moment-matching trials of 0, 50, 75, or 100% MVC. Trials were separated by at least 90 seconds of rest to mitigate fatigue. A supramaximal (i.e. 120 % RMT) TMS pulse was delivered when the participant achieved and maintained ± 2.5 percent of the target effort level for a sustained 0.5s.

### Data Analysis

The peripheral and cortical superimposed twitch (SIT) moments were computed for each trial as the difference between the maximum moment occurring within 150 ms after the stimulus event and the pre-stimulus moment. The pre-stimulus moment was computed as the maximal 10 ms moving average moment maintained within 50 ms prior to the stimulus event. The potentiated resting twitch moment was also computed for each motor point stimulation trial. Peripheral VA was calculated using Equation 1:

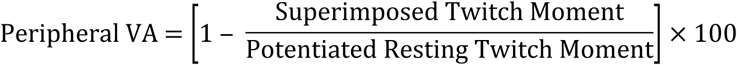

To compute cortical VA, the resting twitch was estimated via linear regression using the methodology described by Todd et al. (2003). The linear regression function was derived from the amplitude of the cortical SIT and the corresponding voluntary elbow moments at 50, 75, and 100% MVC (Figure 1). Cortical VA was calculated using Equation 2:

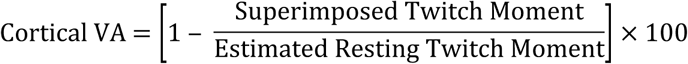

Cortical VA blocks with poor linearity of voluntary torque and SIT were excluded (r < 0.8); 7 out of 50 blocks were excluded for this reason. All MEPs recorded during a given session were normalized to the Mmax (biceps or triceps as appropriate) value for the corresponding session (18,19). The biceps/triceps MEP ratio for each trial was then calculated as the normalized biceps MEP divided by the normalized triceps MEP. To determine the effect of the MEP ratio on cortical VA, MEP ratios were averaged across effort levels to represent its overall magnitude throughout any given block for each participant.

### Statistical Analysis

A linear mixed effect model was analyzed to determine the effect of independent variables on cortical VA (the dependent variable). The independent variables were: isometric elbow flexion angle, block mean biceps/triceps MEP ratio, linearity of the voluntary torque and superimposed twitch relation (r value), and RMT. In addition to excluding blocks with low linearity (r < 0.8), we included linearity as an independent variable to test whether smaller variations in linearity affect the estimation of cortical VA. Resting motor thresholds were added to the model as a continuous covariate. A random effect was added to account for individual differences that resulted in each participant being assigned a different intercept. P-values were obtained via the Kenward-Roger approximation for degrees-of-freedom implemented for linear mixed effect models. Comparisons were reported with respect to 90° elbow flexion, which is the common elbow angle used in previous studies investigating cortical VA of the elbow flexors (6,20).

A two-way ANOVA with repeated measures and a Fisher’s least significant difference post-hoc test were used to compare the biceps/triceps MEP ratio across elbow angles for each target effort level. Another two-way ANOVA was analyzed to determine the effect of elbow angle on normalized MEP amplitudes of the biceps and triceps separately. Intraclass correlation coefficients (21) were determined to assess the intersession repeatability of peripheral and cortical VA. A two-way mixed model, ICC(3,k) was used where sessions are considered fixed effects and participants were treated as random effects. The mean of k measures was used with k = 10 for peripheral VA, and k = 6 for cortical VA. Coefficients of variation (SD/mean) were computed per participant and per session then averaged to represent within-session variability of cortical VA measures. All data and statistical analyses were performed in Matlab (MathWorks, Inc, Natick, MA), R (R Core Team, Vienna, Austria) and Prism (GraphPad Software, La Jolla California USA) with custom-written code. Tests were evaluated at a significance level corresponding to p < 0.05.

## RESULTS

Across all participants, mean cortical VA at 90° elbow flexion was 92.1%, 94.6%, and 93.8% for sessions 1, 2, and 3, respectively. Mean cortical VA was 90.3% at 120° elbow flexion, and 89.6% at 45° elbow flexion. Mean peripheral VA was 98.0%, 95.9%, and 97.8% for sessions 1, 2, and 3, respectively. A summary of all key measures collected is presented in Table 1.

**Table 1:**
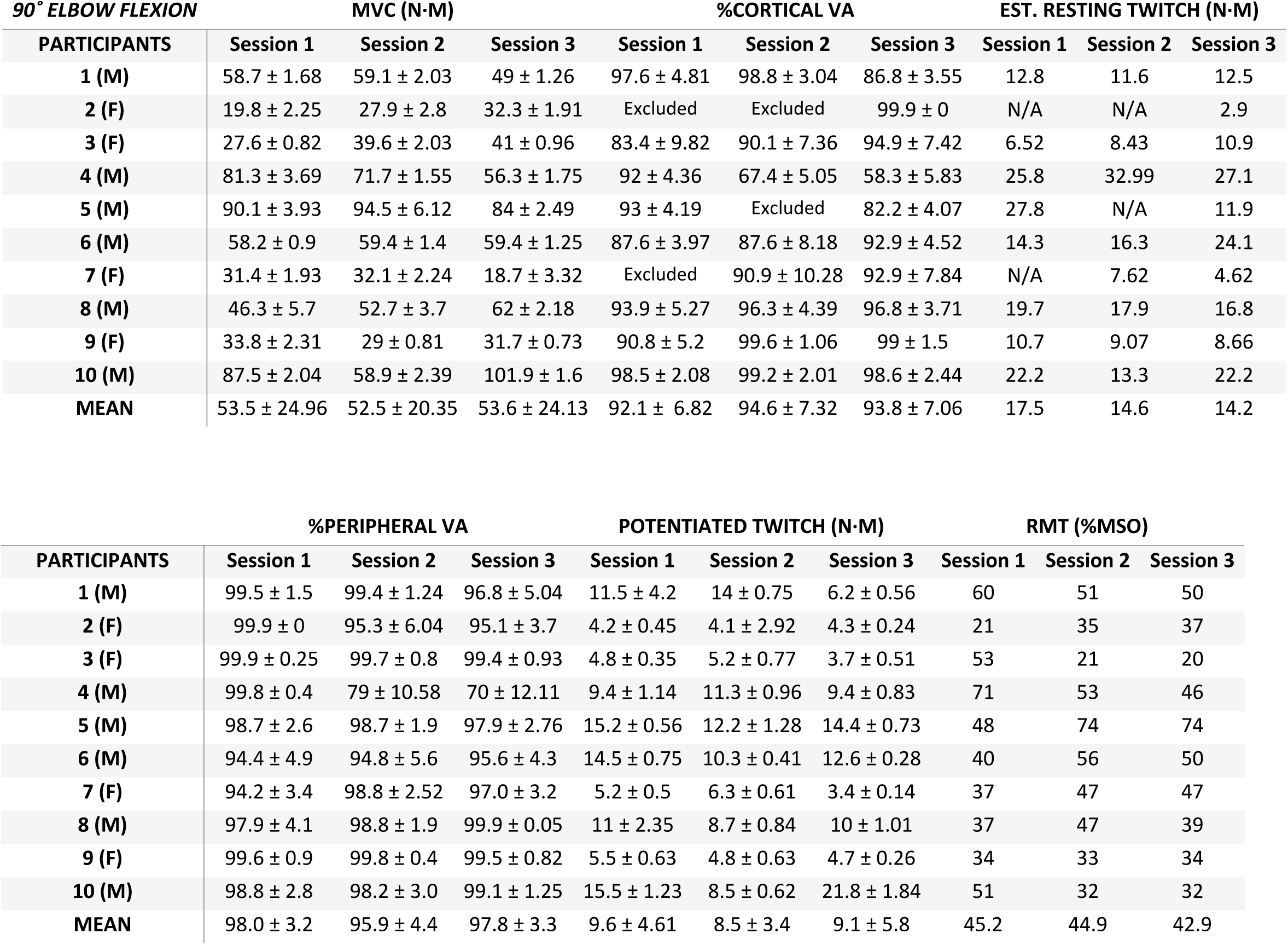
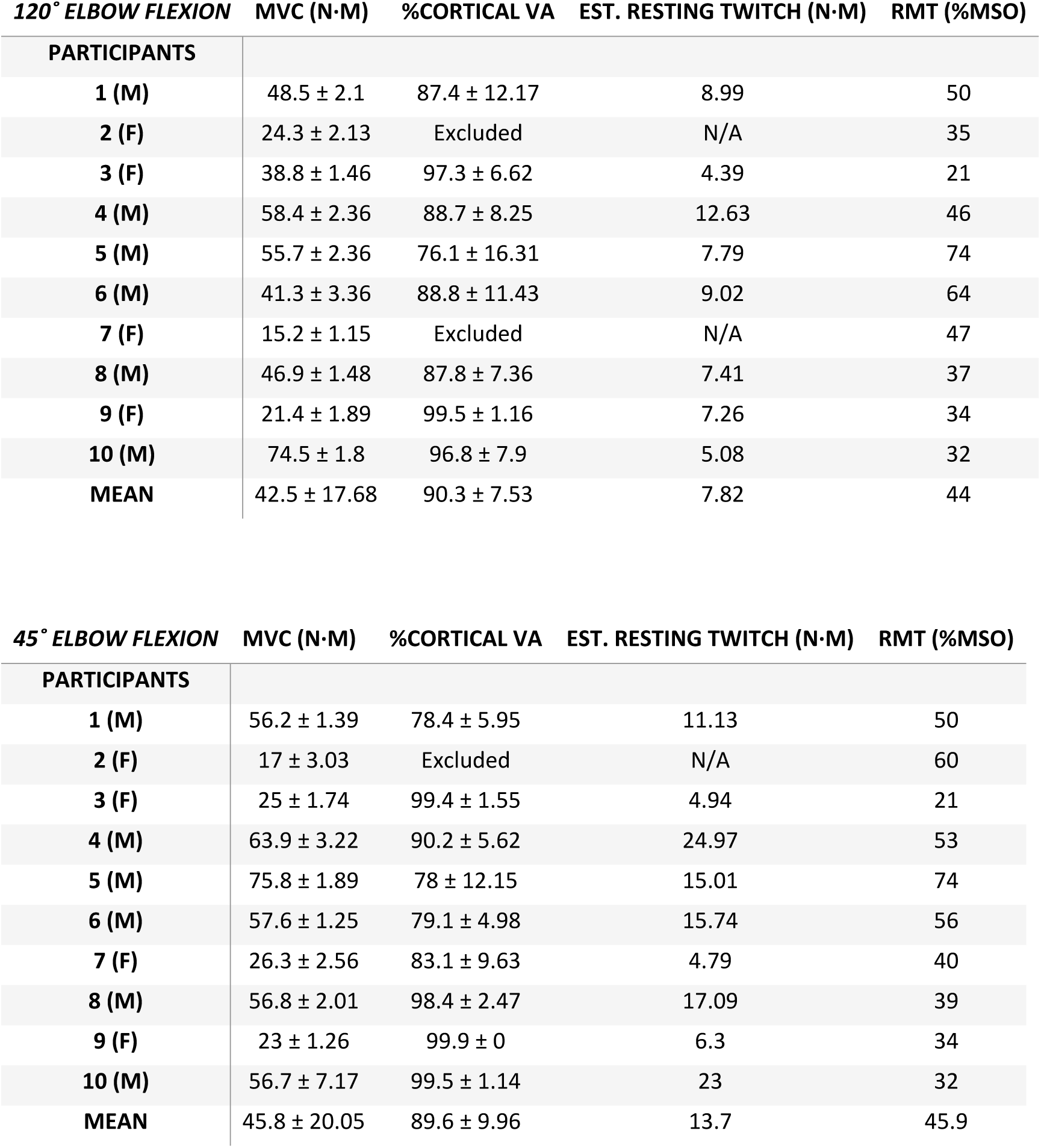
Data summary presenting all the key measures collected and calculated during the study. When applicable, measures are presented as mean ± standard deviation.

### Effect of Elbow Angle on the Biceps/Triceps MEP Ratio

Across all elbow angles, the biceps/triceps MEP ratio increased from 0 to 50% MVC, then decreased from 50% to 100% MVC [p < 0.0001]. At rest (i.e., 0% MVC), there were no differences in the biceps/triceps MEP ratio due to the elbow angle. At 50% MVC, the biceps/triceps MEP ratio was greater in 120° flexion relative to 90° flexion [Figure 2, p = 0.033]. At 75% MVC, biceps/triceps MEP ratio in 120° was greater relative to 90° flexion [p = 0.009]. At 100% MVC, biceps/triceps MEP ratios did not differ across the elbow angles (Figure 2).

**Figure 2.**
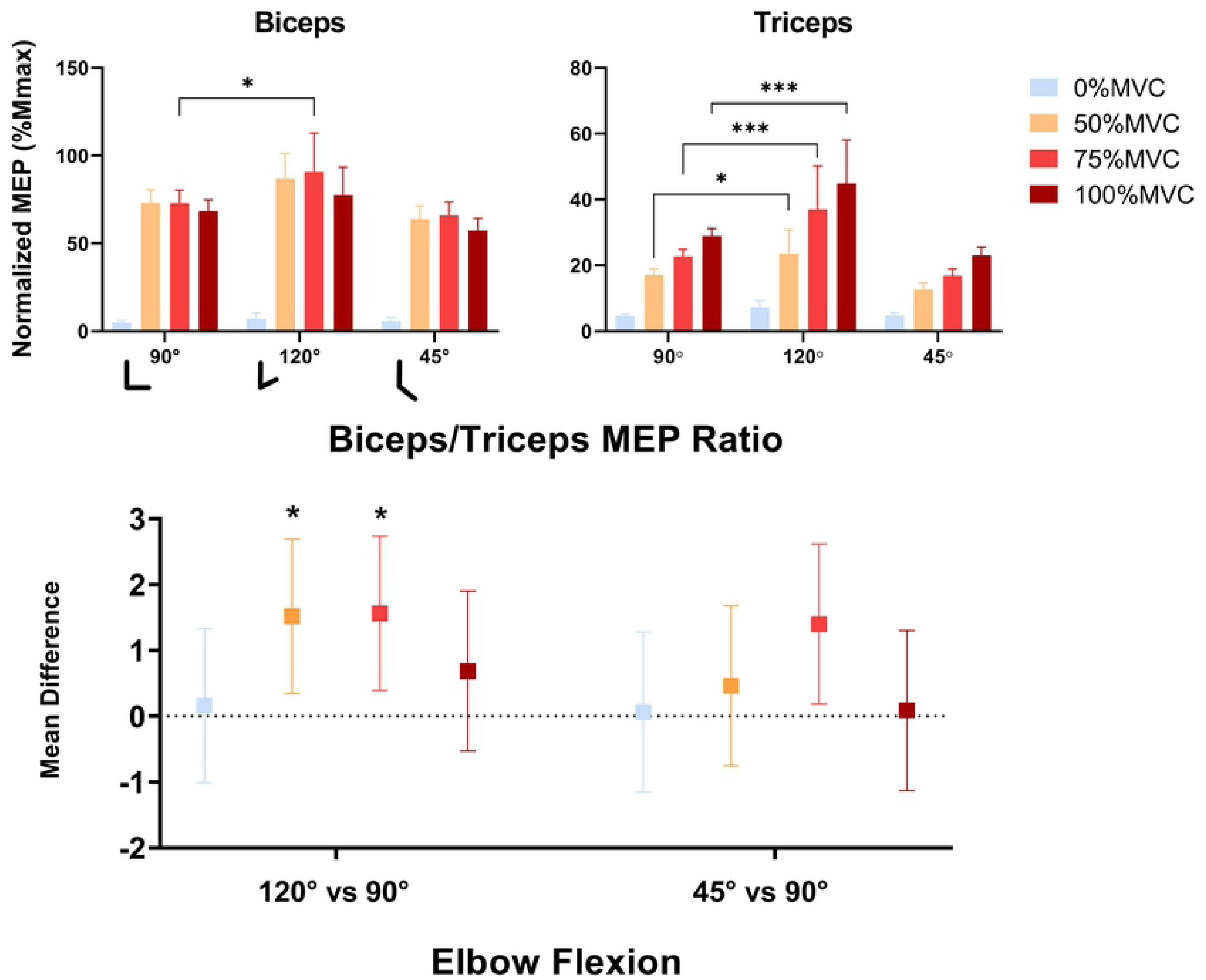
Top: Average biceps and triceps MEPs (normalized to corresponding Mmax) across elbow angles and effort levels. Biceps MEPs increased significantly from rest to 50% MVC then reached a plateau while triceps MEPs increased linearly with effort. Error bars represent 95% confidence intervals. Bottom: MEP ratio mean difference relative to 90° of elbow flexion across effort levels. Errors bars show 95% confidence intervals and asterisks indicate a significantly greater mean MEP ratio (p < 0.05) relative to the mean MEP ratio at 90° elbow flexion.

### Effect of Independent Variables on Cortical VA

The main effect of elbow angle on cortical VA and the interaction effect of elbow angle and biceps/triceps MEP ratio on cortical VA were significant in the linear mixed-effects model. Cortical VA assessed at an isometric elbow angle of 45° flexion (i.e., the more extended elbow angle) was 14.6 ± 4.2% lower relative to cortical VA assessed in 90° elbow flexion [t = −3.46, p = 0.0019]. For each unit increase in the biceps/triceps MEP ratio, cortical VA at 45° elbow flexion is predicted to increase by 2.84% [t = 3.08, p = 0.0035]. The fixed effect of RMT on cortical VA was significant. The model predicted that cortical VA would decrease 1.82% for each 10% maximum stimulator output (MSO) increase in RMT [t = 2.39, p = 0.033]. Interaction analyses showed that this effect was independent of the elbow angle, linearity, and MEP ratio. The fixed effect of the linearity of the voluntary torque and SIT relation on cortical VA was significant. For each 0.1 increase in linearity (for 0.8 < r < 0.99), cortical VA was predicted to increase by 4.4% [t = 2.34, p = 0.023]. This effect was independent of the elbow angle, MEP ratio, and RMT.

### Repeatability and Variability of VA Estimates

An ICC of 0.64 [p < 0.01] resulted from the inter session analysis of peripheral VA assessed in 90° elbow flexion. An ICC of 0.71 [p < 0.01] resulted from the inter session analysis of cortical VA assessed in 90° elbow flexion. (Figure 3). Mean within-session coefficients of variation for cortical VA were 6.2 ± 5% at 90° elbow flexion, 14.7 ± 10.5% at 120° elbow flexion, and 5.8 ± 4.9 % at 45° elbow flexion.

**Figure 3.**
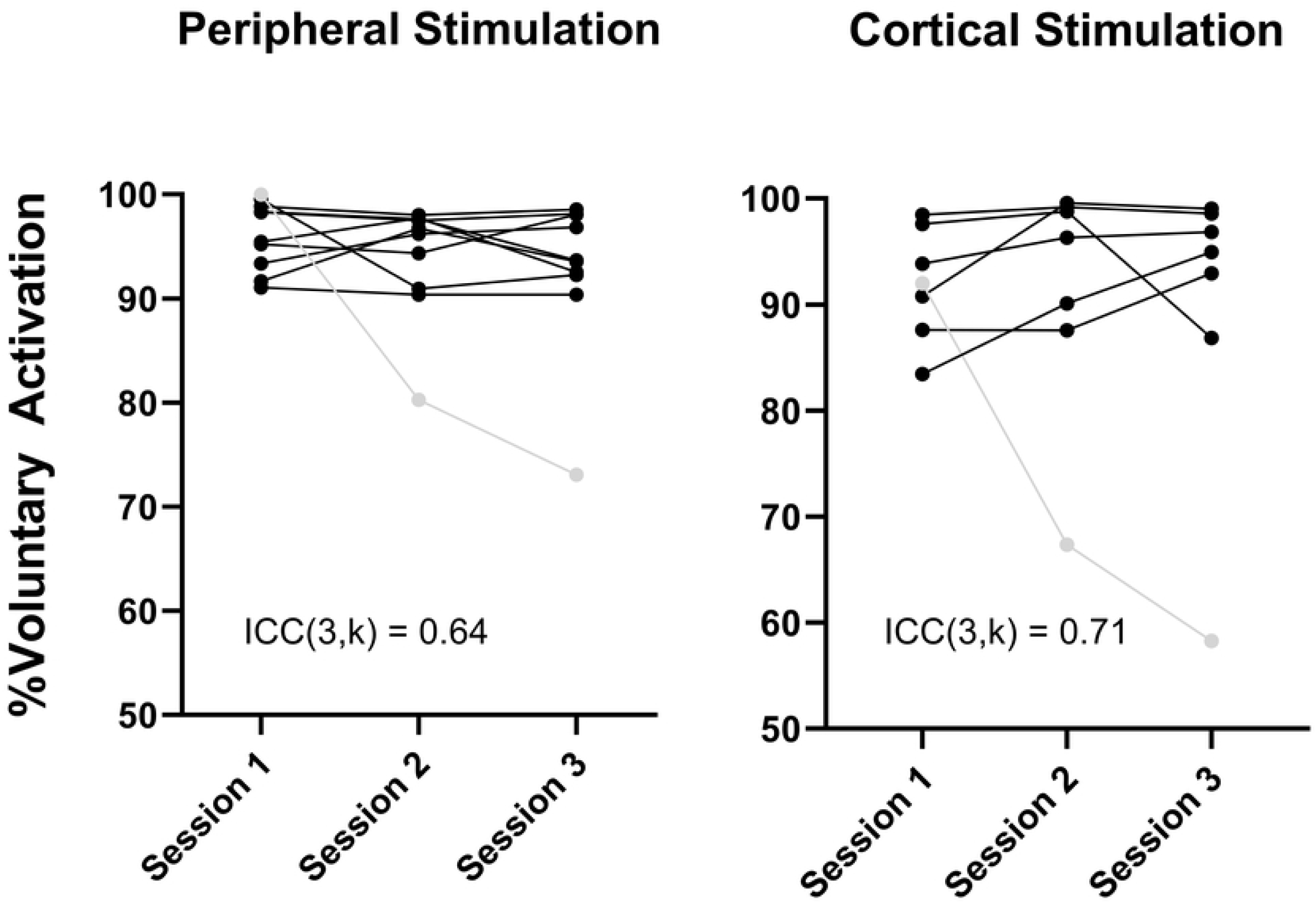
Interclass correlation coefficients (ICC) suggest that peripheral (left) and cortical (right) VA are repeatable at 90° of elbow flexion across three sessions. Each data point represents the mean VA estimate for a participant and session. The participant depicted in light gray was determined to be abnormally fatigued in sessions 2 and 3. The participant’s data were subsequently excluded from the analysis.

### Post-hoc Evaluation of Biceps/triceps MEP Ratio

Post-hoc evaluation was performed to further analyze the biceps/triceps MEP ratio. We determined how often the recommended cortical stimulation condition was achieved (7). The percent of trials with a biceps/triceps MEP ratio above or equal to 2.5 for each elbow flexion angle and across all participants is presented in Table 2.

**Table 2:** Percent of cortical stimulation trials that produced a biceps/triceps MEP ratio greater or equal to 2.5 for each elbow angle and effort level.

| Effort levels | Isometric Elbow Flexion Angle |  |  |
| --- | --- | --- | --- |
|  | 90° | 120° | 45° |
| 50% MVC | 77 | 89 | 94 |
| 75% MVC | 61 | 83 | 72 |
| 100% MVC | 46 | 54 | 61 |

We tested for correlations between RMT and the biceps/triceps MEP ratio. Resting motor thresholds were negatively correlated with the biceps/triceps MEP ratio at 0% MVC only [r = – 0.34, p < 0.0001] (Figure 4). At rest, a low negative correlation was present [r = −0.34, p < 0.0001] and while this trend continued during effort, the correlation was weak [at 50% MVC: r = −0.17, p = 0.0002; at 75% MVC: r = −0.23, p < 0.0001; at 100% MVC: r = −0.16, p = 0.0003].

**Figure 4.**
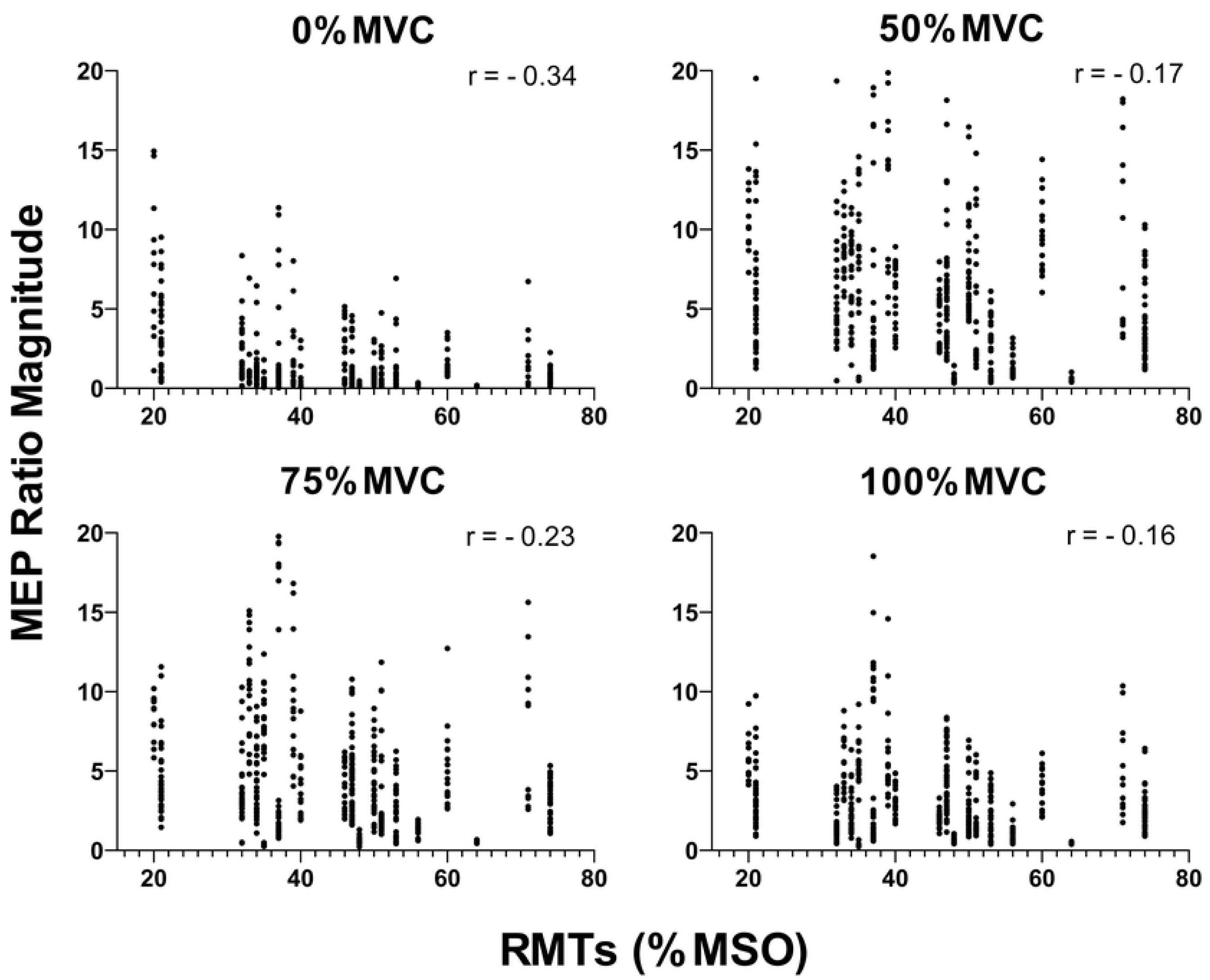
Correlation between MEP ratios (%Mmax) and RMTs across effort levels. MEP ratio and RMTs were negatively correlated during the resting condition only (0% MVC, top left). A total of 492 matching pairs (participant and session matched) per effort level were included in the analysis.

### Effect of Elbow Angle on Biceps MEPs

At 0% MVC, there were no differences in biceps MEPs due to elbow angle. At 75% MVC, 120° of elbow flexion increased normalized biceps MEPs [+11.3% Mmax, p = 0.04] compared to 90° elbow flexion (Figure 2).

### Effect of Elbow Angle on Triceps MEPs

At 0% MVC, there were no differences in triceps MEPs due to elbow angle. At 50% MVC, 120° of elbow flexion increased normalized triceps MEPs [+9.8% Mmax, p = 0.01] compared to 90°. Similarly, at 75% MVC, 120° of elbow flexion increased normalized triceps MEPs [+18.4% Mmax, p < 0.0001] compared to 90°. At 100% MVC, 120° of elbow flexion increased normalized triceps MEP amplitudes [+18.1% Mmax, p < 0.0001] compared to 90° of elbow flexion (Figure 2).

## DISCUSSION

The objectives of this study were to determine the effect of elbow angle on: 1) the biceps/triceps MEP ratio across a range of voluntary efforts, and 2) estimates of biceps cortical VA in non-impaired participants. We hypothesized that the biceps/triceps MEP ratio would be greatest in a more flexed elbow angle (120° flexion) at each level of voluntary effort. This hypothesis was only supported at the 50% and 75% MVC effort levels (i.e., not at 100% MVC). Further, we hypothesized cortical VA would depend on the biceps/triceps MEP ratio. This hypothesis was only supported in the more extended elbow angle (45° flexion). Our results indicate that while the bicep/triceps MEP ratio was modulated by the elbow angle, modulation did not occur across the full range of voluntary efforts and may not improve the estimation of cortical VA. Finally, both cortical and peripheral VA measured at 90° elbow flexion were repeatable across three days.

The more extended elbow angle led to lower cortical VA measures compared to 90° elbow flexion. and this effect was dependent on the mean biceps/triceps MEP ratio (for each given block). In the more extended arm posture, a large biceps/triceps MEP ratio was associated with a greater cortical VA estimate. This effect suggests a small effect of better targeting the biceps relative to the triceps with TMS, but considering this effect was only seen in one condition, other factors effecting cortical VA estimation are also important. The decreased cortical VA in the more extended elbow angle is not due to changes in the moment-generating capacity of the biceps and triceps with elbow angle because VA is expressed as a ratio of a superimposed torque response to TMS over the estimated resting torque response to TMS. Thus, VA is normalized by the moment-generating capacity at a given elbow angle.

Although the biceps/triceps MEP ratio may reflect the focality of cortical stimulation when targeting the biceps, its contribution to improving the estimation of cortical VA is limited. In order for the biceps/triceps MEP ratio to improve cortical VA estimation, the ratio must be increased across the range of effort levels needed to assess cortical VA (i.e., 50, 75 and 100% MVC). In the more flexed elbow angle (120°), we observed increased MEP ratios at only 50% and 75% MVC. Furthermore, the increased MEP ratio in the more flexed elbow was not associated with a change in the magnitude of cortical VA. The increased biceps/triceps MEP ratio in the more flexed posture occurred primarily via biceps MEP facilitation (Figure 2). Changes in MEP amplitudes due to isometric joint angle are mostly attributed to spinal mechanisms (13,22), namely the influence of afferent feedback provided to the spinal cord (11). However, central influence also plays a role (12). In this study, we modulated the elbow angle while keeping other posture-related parameters (i.e. forearm orientation, shoulder and head position) unchanged. Therefore, the increased biceps MEP ratio is likely due to a combination of central and spinal facilitatory mechanisms affecting the shortened biceps. Additionally, triceps MEP facilitation is higher at 100% MVC compared to 50-75% MVC whereas biceps responsiveness to TMS reaches its peak early, at high but submaximal voluntary contractions, then deteriorates at near-tetanic state of the muscle (23). This may explain why we could detect an increase in MEP ratio only at high but submaximal biceps recruitment.

The effect of RMTs on cortical VA estimation could not be explained by biceps/triceps MEP ratio modulation. We found that higher RMTs were associated with lower cortical VA estimates (although the effect was small: −1.82% cortical VA for each 10% MSO increase). Further analysis showed that RMTs were negatively correlated with the magnitude of the biceps/triceps MEP ratio which may indicate better ability to target the biceps relative to the triceps in individuals with low RMT. Lower stimulus intensities will typically decrease stimulus spread to cortical representations of other muscles (24). However, as this correlation became negligible during voluntary effort, MEP modulation was not involved in the mechanism by which higher RMTs led to lower cortical VA.

The repeatability of cortical and peripheral VA measured at 90° across three sessions was consistent with previous reports suggesting good reliability of these measures (25,26). Inter-session reliability of cortical VA had previously been established across two days in the elbow flexors (20), wrist extensors (27) and knees extensors (28). Peripheral VA assessed in the elbow flexors is reliable across five days (29). Furthermore, while within-session variability was small for cortical VA at 45° (CV = 5.8 ± 4.9 %) and 90° (CV = 6.2 ± 5%) of elbow flexion and peripheral VA, the more flexed elbow flexion presented higher variability (CV = 14.7 ± 10.5%). Reliability of VA baseline measures (i.e. pre-fatigue and/or pre-therapy) is critical for use in clinical settings to monitor patients and to compare cortical and peripheral VA.

Our measures of cortical VA were underestimated compared to peripheral VA. Underestimation of cortical VA compared to peripheral VA has been reported in both unfatigued and fatigued biceps (4,6) and remains a methodological challenge when estimating cortical VA (7). Poor linearity of the voluntary torque and SIT relation may explain artificially low estimation of cortical VA. In this study, modulating the elbow angle had no impact on linearity. Previously, neuromuscular fatigue was associated with a decrease in linearity of the voluntary torque and SIT relation (6). This suggests that fatigue may set in the neuromuscular system in a non-linear way across the range of voluntary effort. As the method used to estimate cortical VA relies on the assumption of linearity between voluntary torque and the SIT, the validity of what is measured may be hindered in a context that affects linearity, such as fatigue. Overall, our analyses further support this claim as we identified a positive association between linearity and the magnitude of cortical VA, even amongst blocks with satisfactory linearity (r > 0.8). Together, this implies that cortical VA is sensitive to the linearity of the voluntary torque and SIT relation, even when linearity is high (0.8< r < 0.99).

A potential limitation of our approach is that we excluded blocks from the data analysis demonstrating poor linearity (r < 0.8). Although excluding data because of poor linearity is a common approach (7,30,31), our exclusion rate was higher (14% versus 7-9%) which may be related to our experimental design with longer blocks, including resting periods between each trial (one per effort level). However, our increased rest period was likely beneficial in preventing fatigue. Only one participant expressed outstanding levels of fatigue (abnormally low MVC, low peripheral and cortical VA) in session 2 and 3 compared to session 1 (Figure 3, light gray). His data were excluded in order to prevent the effects of fatigue from contaminating our analysis. Another potential limitation of using TMS to measure cortical VA is the high variability of MEPs (32) that depends on the levels of muscle activation (33). MEPs indicate how the corticomotor pathway responds to cortical stimuli. Thus, MEP variability may translate to SIT moment variability, which subsequently affects cortical VA. A third limitation is that we normalized all MEPs to the Mmax collected at the start of a session to allow comparison of MEPs between participants and across days. However, Mmax can decrease over the course of a session despite constant isometric posture (34,35) and is sensitive to the level of voluntary contraction (36). Here, we collected Mmax at rest while MEPs were collected at various levels of muscle activation.

In conclusion, our results indicate that modulating the elbow flexion-extension angle may not improve cortical VA estimation in the non-impaired biceps. While the biceps/triceps MEP ratio was modulated by the elbow angle, MEP modulation did not occur across the full range of voluntary efforts and did not uniformly impact cortical VA. Thus, a focus on increasing the biceps/triceps MEP ratio through modulation of the elbow angle does not further improve estimation of cortical VA in non-impaired individuals. Secondarily, cortical VA was sensitive to elbow flexion angle which suggests that elbow angle should be carefully monitored and reproduced across trials, participants, and sessions when assessing VA. Finally, cortical VA was sensitive to small changes in the linearity of the voluntary torque and SIT relation, which plays a role in the underestimation of cortical VA relative to peripheral VA. As modulating elbow flexion did not affect linearity of the voluntary torque and SIT relation, further research is needed to find novel approaches to improve linearity.

## ACKNOWLEDGMENTS

We would like to thank Joshua Arenas for his assistance with the experimental setup and subject recruitment. This work was supported by the Virginia Commonwealth University’s CTSA (UL1TR000058 from the National Center for Advancing Translational Sciences) and the CCTR Endowment Fund of Virginia Commonwealth University. The funders had no role in study design, data collection and analysis, decision to publish, or preparation of the manuscript.

